# Mineral Phase Changes During Intervertebral Disc Degeneration

**DOI:** 10.1101/2024.07.08.602462

**Authors:** Theresa Banu Yenen, Ravin Jugdaohsingh, William D. Thom, Sam Khan, Viviana Rojas Solano, Giulio Lampronti, Andy Brown, Janire Saez, Davide Corbetta, Salih Eminağa, Giunio Bruto Cherubini, Jonathan Powell, Kate Hughes, Paul Freeman

**Author notes:** corresponding author Paul Freeman.

## Abstract

Intervertebral disc disease is a common cause of pain and neurological deficits and is known to be associated with degeneration and calcification. Here we analysed samples of herniated disc material and compared it to material taken from non-herniated discs following surgical treatment in dogs. Our clinical approach to these cases allows collection of samples providing a unique opportunity for a case-controlled study such as this, an opportunity which is not available to the human neurosurgeon. We analysed all samples using Fourier transform infrared (FTIR) spectroscopy, as well as a proportion with X-ray diffraction (XRD) and transmission electron microscopy (TEM). FTIR spectra of the majority of herniated samples were consistent with the presence of crystalline hydroxyapatite, whereas most of the non-herniated discs showed spectra consistent with amorphous phosphate material. XRD analysis and TEM confirmed these findings and identified the amorphous material as amorphous calcium phosphate nanoparticle clusters of ∼ 20 nm diameter and the crystalline hydroxyapatite material as needles up to 100 nm in length.

The differences between the herniated and non-herniated discs indicate that the degenerative process involves a conversion of amorphous calcium phosphate into crystalline hydroxyapatite which precedes and may predispose the disc to herniate.

## INTRODUCTION

Intervertebral disc disease is characterized by the degeneration of disc(s) that separate the bones of the spine leading to localised pain that frequently extends to the legs and arms. It affects ∼ 5% of the population in high income countries and costs the UK National Health Service an estimated £5 billion p.a. (source Manchester Metropolitan University). Moreover, the disease is not restricted to humans. Chondrodystrophic dogs, characterised by a short-legged conformation, such as the dachshund and French bulldog, are especially prone to frequent, severe disease.

The intervertebral disc of mammals consists of the inner gelatinous nucleus pulposus, the fibrous annulus fibrosus which creates a border around the nucleus, and the cartilaginous endplates which bind the disc to the adjacent vertebral bodies. The healthy nucleus pulposus contains mainly notochordal cells surrounded by a matrix rich in proteoglycan and collagen (type II) as well as hyaluronic acid, creating a high osmotic pressure leading to a high-water content ^1-6^. Calcification of intervertebral discs, in both the annulus fibrosus and the nucleus pulposus, is seen in disc degeneration of dogs and humans ^7-11^. In dogs, calcification is associated with disc extrusion ^12,13^ which can lead to paralysis and incontinence or even death, whilst in humans it is associated with discogenic pain ^14-16^. However, whilst calcification is well recognised in the pathogenesis of intervertebral disc disorders, the relationship of the mineral composition to clinical disease is not known. At the cellular level, amorphous calcium phosphate particles are benign but ectopic hydroxyapatite crystals are pro-inflammatory so mineral phase could well dictate outcomes in calcified discs ^17^. Indeed, hydroxyapatite has been found in both degenerated ^18,19^ herniated ^20^ human intervertebral discs as well as in joints of patients with osteoarthritis ^21^. In addition, large crystal-sized hydroxyapatite accumulations (80-130 nm) were found in ovine intervertebral discs and were considered to be related to aging ^22^. In contrast, little is known about the presence of amorphous calcium phosphate in intervertebral discs despite many studies supporting the possibility that amorphous calcium phosphate could be the precursor form of hydroxyapatite in other biological situations such as bone formation ^23-26^. During surgery for extruded intervertebral discs in dogs, in addition to removing herniated nucleus pulposus material from the vertebral canal, it is good practice to clear out non-extruded neighbouring discs, a process known as disc fenestration which has been shown to be prophylactic in preventing future extrusion ^27,28^. This provides a unique opportunity to compare mineral phase of the calcified deposits in a clinically degenerative situation with very closely matched non extruded discs, in a way that is not ethically possible in humans. Here we show that the major mineral phase present in extruded canine discs is crystalline hydroxyapatite, whereas a significant proportion of non-extruded discs in the same animals contain amorphous calcium phosphate, providing compelling evidence that a switch from amorphous calcification to crystalline calcification may well precede and trigger degenerative disc disease.

## RESULTS

In total, 57 clinical cases were included, all of which were chondrodystrophic or chondrodystrophic-mix dogs. Clinical details are contained in Supplementary Figure 1. The clinical cases generated 104 individual samples of intervertebral disc material, of which 58 were samples of extruded (herniated) material removed at surgical decompression, and 46 were non-extruded (control) samples of nucleus pulposus taken by fenestration of non-herniated discs.

### FTIR Analysis of Intervertebral Discs

FTIR spectra of 104 disc samples, 58 extruded and 46 non-extruded, were produced. Smoothed spectra examination revealed that a majority of the extruded samples showed a PO_4_^-3^ ν_3_ peak suggestive of significant mineral content. Most of these showed a relatively sharp peak more consistent with crystalline mineral content, although this was not the case in 8 of the samples where the peak was more shallow and broad. 47/58 of these extruded spectra contained an additional PO_4_^-3^ ν_1_ peak. From the non-extruded (control) samples, the PO_4^-3^_ ν_3_ peak was visible in 41/46 of the samples, although sharp and well-defined in just 10, with the PO_4_^-3^ ν_1_ peak present in just 7/46.

Dot plots created from peak maxima values in the FTIR second derivative spectra at predetermined functional group regions revealed differences between extruded and non-extruded disc samples at the mineral diagnostic wavenumbers (**Figure 2**). **Figure 2A** shows that, consistent with the smoothed spectra analysis reported above, both sets of samples showed very similar peak maxima for the ν_3_ PO_4_^-3^ peak indicating the presence of mineral phosphate content in roughly equal proportions between both groups. However, there was a significant difference in the ν_1_ PO_4_^-3^ peaks between extruded and non-extruded samples (**Figure 2B**) indicating the presence of crystalline carbonated-hydroxyapatite in a much greater proportion in the extruded than the non-extruded samples. This difference in intensity was supported by the analysis of the carbonate peak which was again more prevalent in the extruded rather than non-extruded samples (**Figure 2C)**.

**Figure 1.**
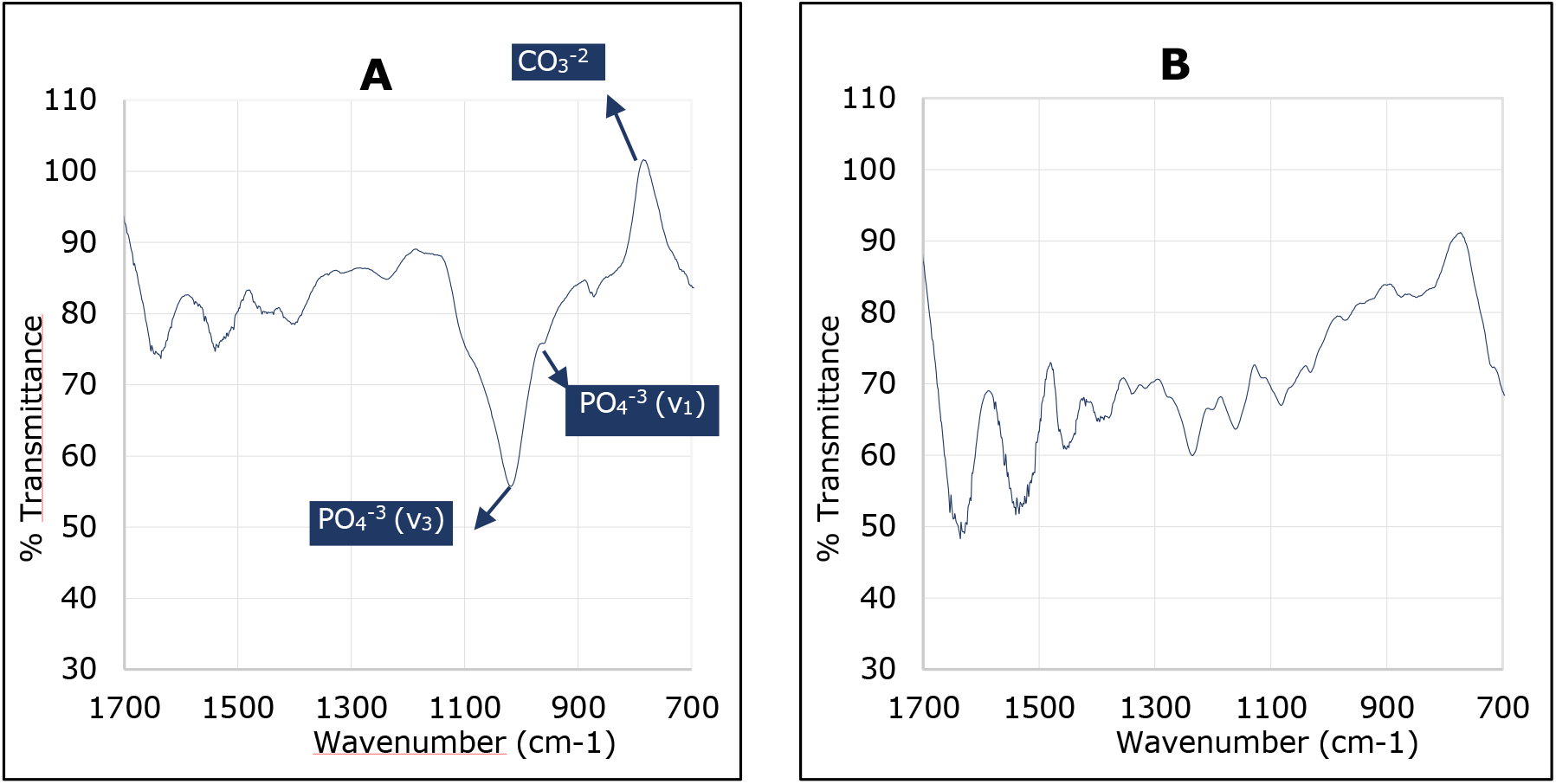
Smoothed FTIR Spectra of Representative Extruded(A) and Non-extruded(B) Intervertebral Disc Materials.

**Figure 2.**
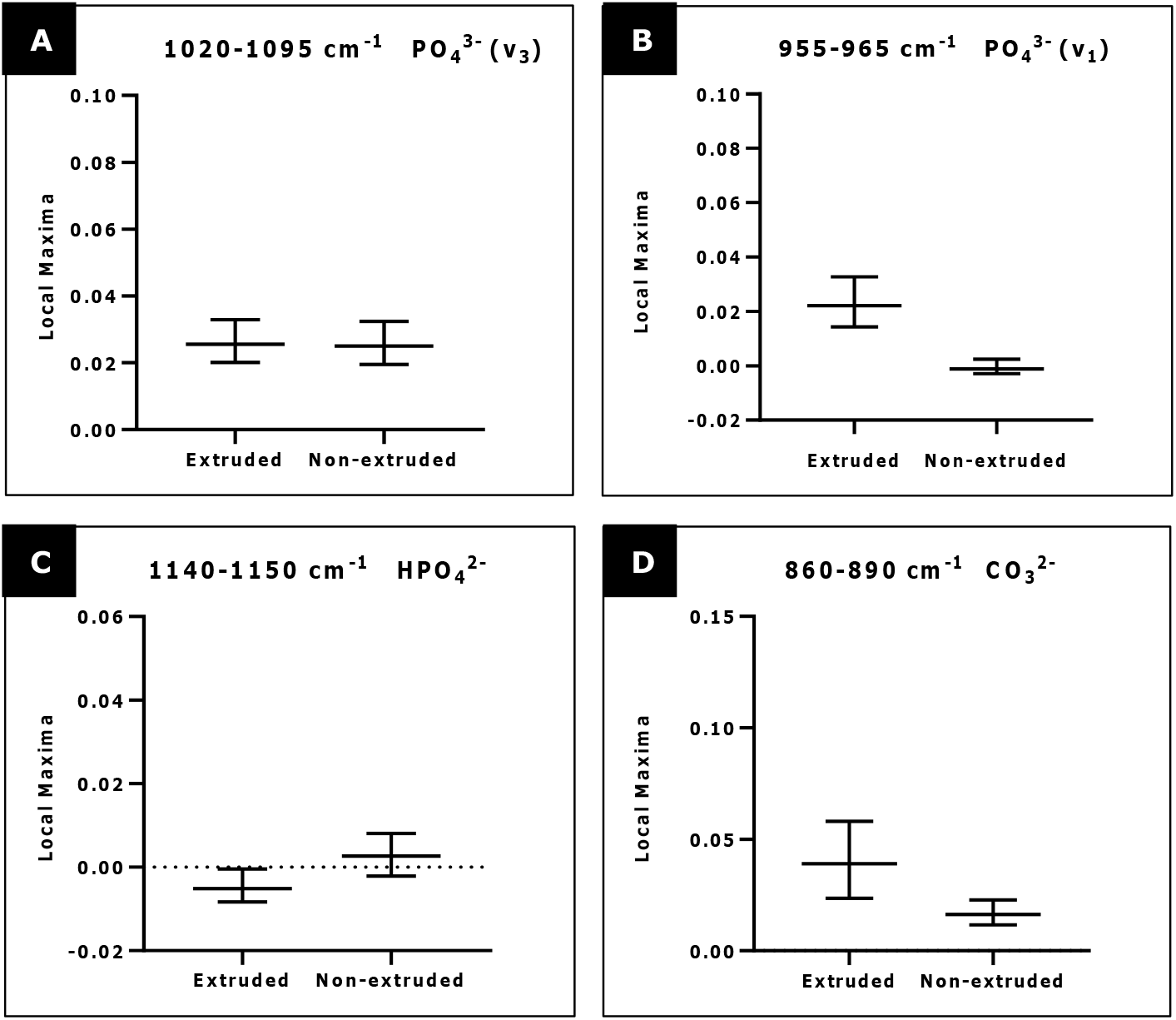
Dot Plots from the Second Derivative Analysis of the FTIR Spectra of IVD Samples (See also Supplementary Figure 3. *Note*. Dotplots showing local maxima values of (A) PO_4_^-3^ v3; (B) PO_4_^-3^ v1; (C) HPO_4_^-2^ and (D) CO_3_^-2^ of extruded (red) and non-extruded (blue) IVD samples

### Powder X-Ray Diffraction Results

The XRD profiles of all five of the extruded (E1-5) and one of the five non-extruded disc materials (N1) were indicative of crystalline hydroxyapatite by the presence of a significant 2θ peak at around 15°, indexed according to the Inorganic Crystal Structure Database ^29^.The other four non-extruded samples (N2-5) showed profiles without any significant peaks indicating that these contained amorphous material only (**Figure 3**).

**Figure 3.**
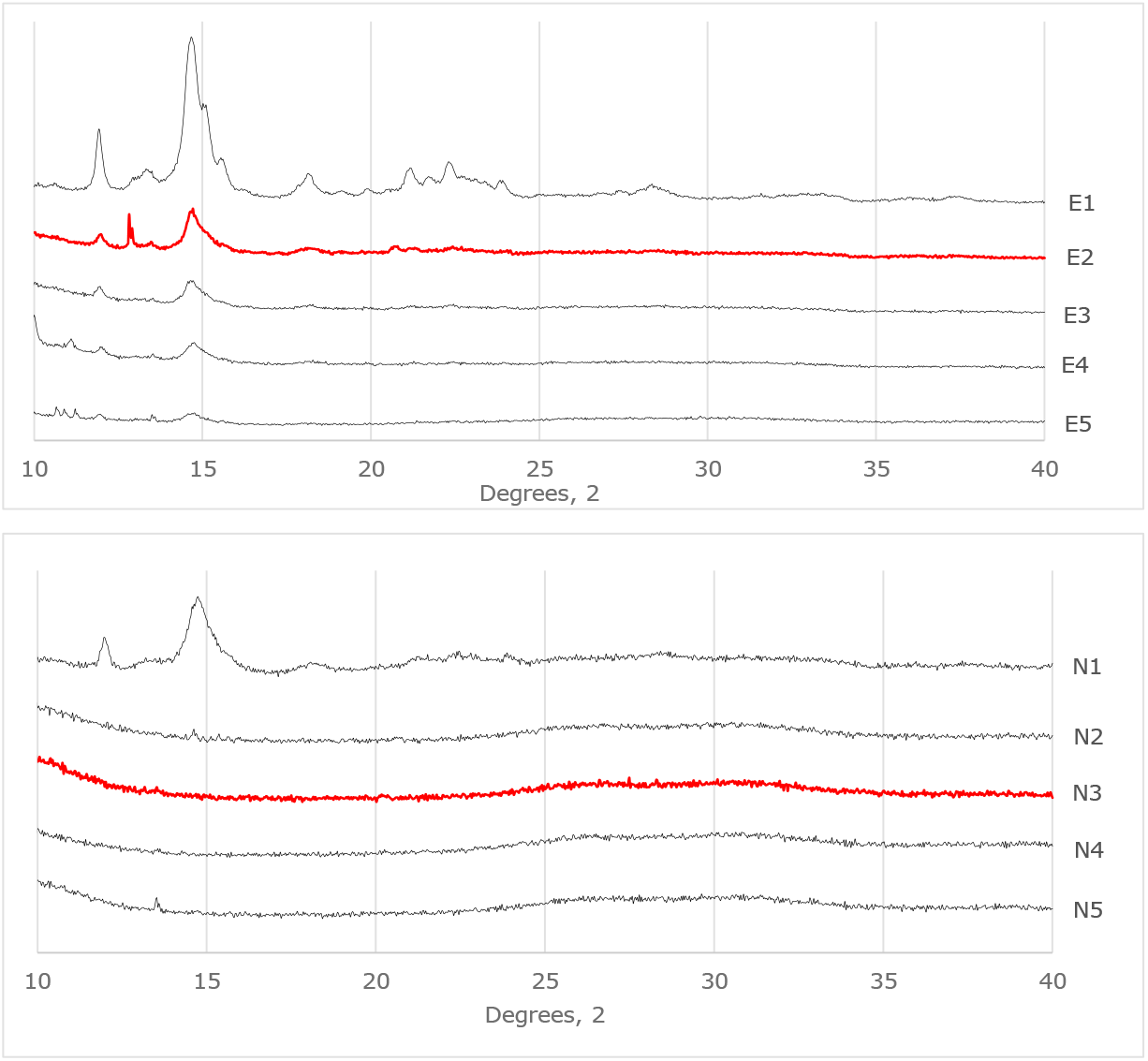
XRD Profiles of 5 Extruded (E1-5) and 5 Non-extruded (N1-5) Samples of Disc Materials. Note. XRD profiles of IVD samples. Samples that were TEM analysed are marked in red and are bold. N: non-extruded; E: extruded.

Crystal sizes of the crystalline hydroxyapatite mineral ranged from 5-15 nm (estimated by peak broadening analysis using the Scherrer equation). The crystal sizes of the hydroxyapatite agglomerates were confirmed by XRD line-broadening analysis and although results varied particle sizes in 4/5 of the extruded samples were notably larger (ranging from 12-15 nm) when compared to the crystal size in the one non-extruded sample which contained hydroxyapatite (8 nm). Details of the Rietveld analysis are reported in the supplementary information.

Dot plots of the maxima intensity values at the same mineral diagnostic wavenumbers as used in Figure 3 following FTIR analysis of these ten samples were created. Among these samples, the 6 hydroxyapatite-containing samples (E1-5 and N1) had higher PO_4_^3−^ ν_1_ maxima values than the 4 X-ray amorphous samples (N1-4) (**Figure 4**).

**Figure 4.**
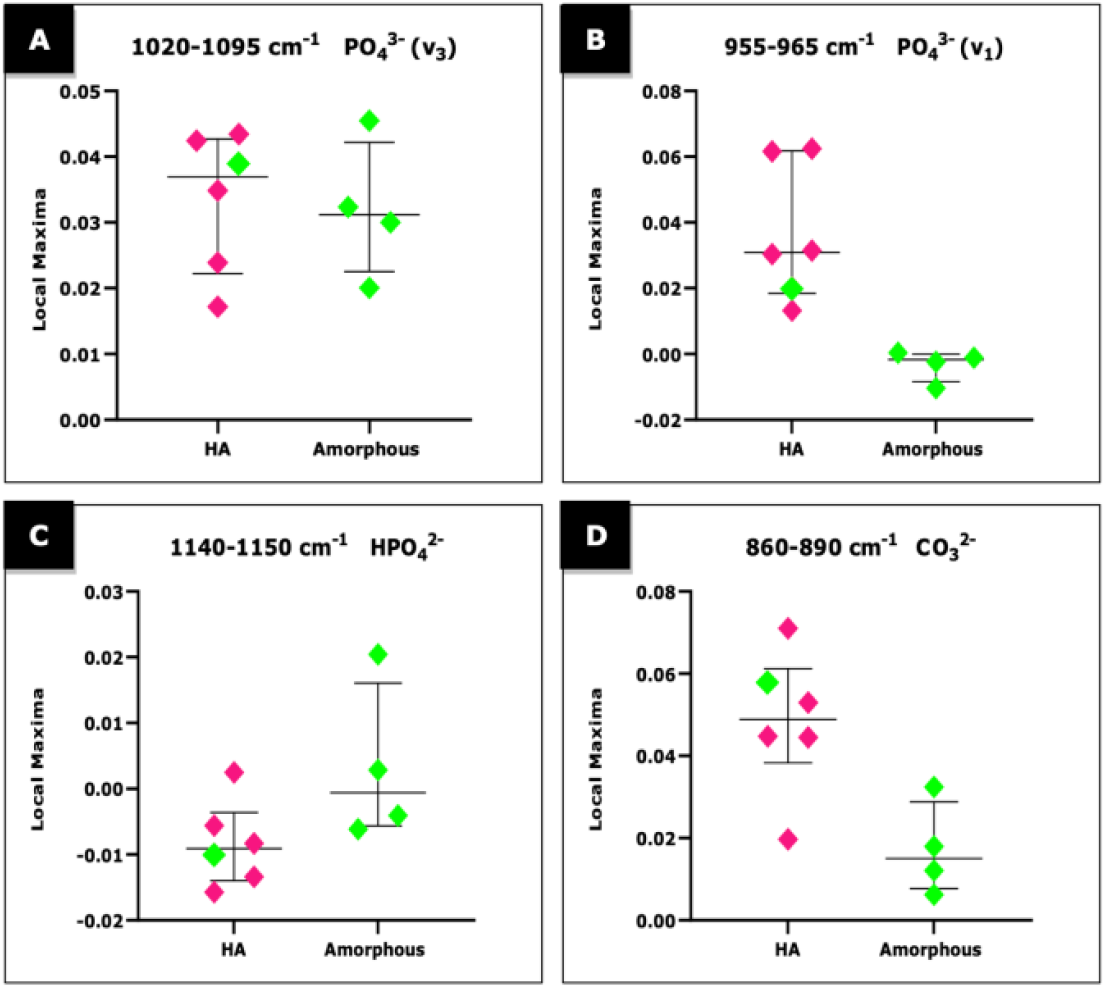
Dotplots from the Second Derivative Analysis of the IVD Sample’s analysed with XRD. Note. Dotplots showing local maxima values of (A) PO_4_^-3^ v3; (B) PO_4_^-3^ v1; (C) HPO_4_^-2^ and (D) CO_3_^-2^ of extruded (pink) and non-extruded (green) IVD samples

### Electron Microscopy Analysis of Two Representative Samples

TEM analysis of one extruded (E2) and one non-extruded (N3) intervertebral disc sample were supportive of both the FTIR and XRD findings. STEM images of the extruded disc material revealed a continuous, gel-like or amorphous phase containing many nanoscale, needle-like agglomerates or deposits within the matrix (**Figure 5A**; the crystalline needles scatter more strongly than the amorphous matrix and so have a white contrast in the HAADF-images. (See also **Supplementary Fig 6A and 6B**). STEM-EDX elemental analysis indicated that the needle–like agglomerates were identified to be Ca, P and O rich (**Supplementary Figure 5**). Furthermore, electron diffraction of the extruded sample showed four polycrystalline rings of spacing consistent with those expected for hydroxyapatite ^30^ (**Figure 6A**).

**Figure 5.**
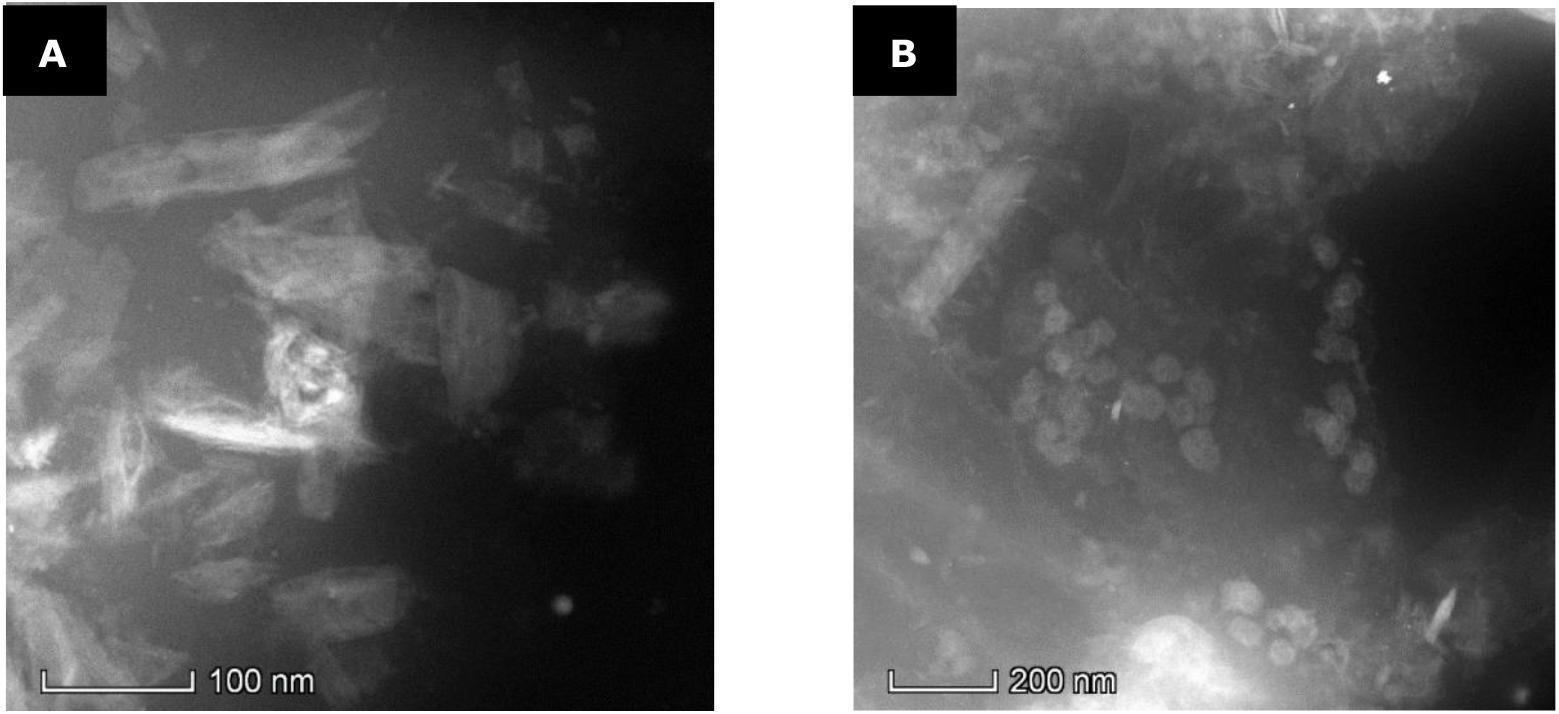
HAADF STEM Images of the Extruded (A) and Non-extruded (B) Intervertebral Disc Material. *Note*. (A) HAADF-STEM of the extruded disc material showing agglomerates of needles indicative of hydroxyapatite nanocrystals; (B) HAADF-STEM of the non-extruded disc material containing small amorphous spheres and only some needle-like particles

**Figure 6.**
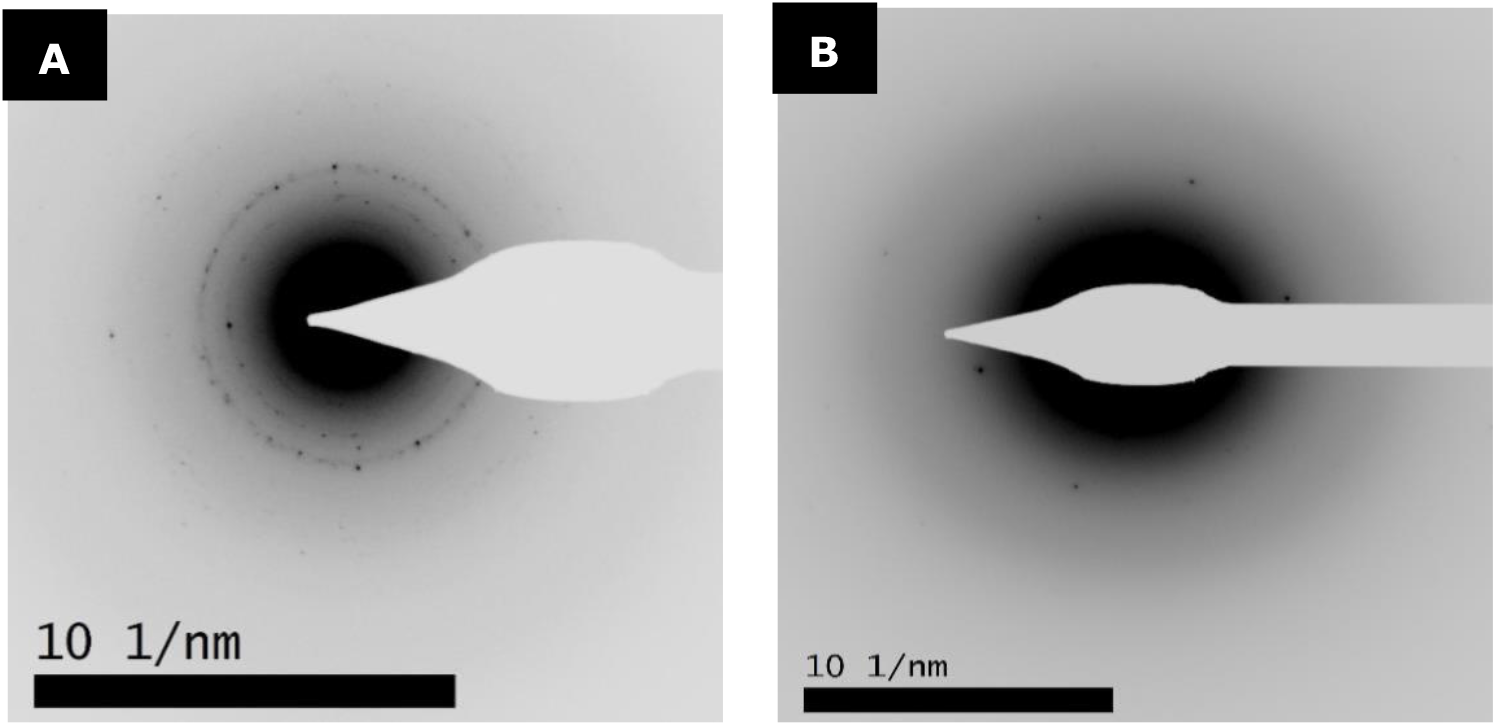
Electron Diffraction Pattern of Extruded (A) and Non-extruded (B) Intervertebral Disc Material. Note. (A) Electron diffraction pattern of extruded IVD material showing four rings at 2.82 (strong ring), 3.2, 3.5 and ∼ 4.4 Å of which the ring spacings are indicative of hydroxyapatite; (B) Electron diffraction pattern of non-extruded IVD material showing diffuse rings with very few diffraction spots that indicate the tested material is predominately amorphous.

TEM images of the non-extruded disc material revealed gel-like deposits similar to the extruded sample. The deposits within the matrix consisted of nanoscale, spherical agglomerates identified as amorphous material by i. the weaker scattering (**Figure 5B**; the deposits are smaller, rounder and less white than the needles in the extruded material in **Figure 5A)** and ii. no visible atomic lattice in high resolution imaging (**Supplementary Figure 6C and D**). In addition, electron diffraction analysis confirmed that the deposits in the gel are predominately amorphous by a lack of sharp rings and spots (**Figure 6B**). The spherical agglomerates in the non-extruded sample that were amorphous by electron diffraction were found to be Ca, P and O rich by EDX, consistent with an amorphous calcium phosphate phase (**Supplementary Figure 7)**.

## Discussion

This study produces new information regarding the mineral composition of calcified degenerate intervertebral discs in chondrodystrophic dogs, revealing that the discs contain either crystalline hydroxyapatite or amorphous calcium phosphate nanoparticles. Amorphous calcium phosphate nanoparticles were found only in discs which had not undergone extrusion (or herniation), whereas crystalline hydroxyapatite was found in both herniated and non-herniated discs. These findings suggest that canine intervertebral disc degeneration, at least in this group of chondrodystrophic dogs, involves a transformation (change of phase) from an initially amorphous calcium phosphate phase into crystalline hydroxyapatite nanoparticles. It is possible that this phase change may then predispose the discs to extrusion.

To minimise the possibility of mineral phase changes post sample collection, freeze-dry**D**ing was used for sample preparation prior to analysis. Freeze-drying has been shown to conserv**D**e the phase and morphology of minerals, although it can cause particle size reduction ^31^. It is also possible that the mineral phase of samples of nucleus pulposus changed following extrusion i.e., once inside the vertebral canal and outside the disc, a possibility which could be tested in the future by comparing nuclear material remaining within an extruded disc with the herniated, extruded material. This however seems unlikely given that some non-extruded samples also contained crystalline material.

In this study all samples were analysed by FTIR and, in a representative sample of these, X-ray diffraction confirmed the findings that amorphous calcium phosphate nanoparticles were found only in discs which had not undergone extrusion, whereas the crystalline hydroxyapatite was found in both extruded and non-extruded discs. A single sample from each group, which from visual evaluation of FTIR spectra appeared typical, were further submitted for transmission electron microscope (TEM) analysis to reveal the nanoscale phase and structure of the investigated materials and to confirm that when crystalline the hydroxyapatite phase forms needles.

A small number of the extruded samples did not reveal any of the prominent phosphate or carbonate peaks consistent with biological hydroxyapatite ^32-35^. Although it is possible that these extruded samples were not calcified or did not include crystalline hydroxyapatite, it is more likely that a sampling error might be the reason for these findings, given the clinical nature of this study.

Crystal sizes in the samples analysed with XRD revealed generally larger crystal agglomerates in the extruded than the single non-extruded sample. The size of the crystalline needles observed by TEM were bigger (50 -100 nm) suggesting these are polycrystalline agglomerates (that could have grown from the smaller amorphous calcium phosphate particles seen in the non-herniated disc material). Previous literature suggests that the size of the crystalline particle of hydroxyapatite in bone increases with maturity ^35, 36, 37^. It is possible that crystal, and agglomerate sizes of the hydroxyapatite within the intervertebral discs increases with the stage of degeneration, and may also be associated with extrusion, but larger sample size analysis is required to make a more definitive statement regarding this.

A few of the non-extruded samples showed FTIR spectra consistent with crystalline hydroxyapatite (and one had an XRD pattern confirming this). This is not unexpected if the degenerative process involves a transformation of amorphous calcium phosphate to crystalline hydroxyapatite prior to extrusion, since it may be that a critical mass and/or size of hydroxyapatite is needed before herniation/extrusion occurs. Clearly it is likely that multiple other factors such as mechanical stress, dehydration and loss of proteoglycans and genetic predisposition are also involved in intervertebral disc disease and herniation^38^. However, if discs containing crystalline hydroxyapatite could be identified before they have herniated eg by calibration to MRI, it may be possible to target them with a prophylactic therapy such as surgical fenestration, chemonucleolysis or laser ablation ^27, 28, 39, 40^ .

This study sheds some light on the process of calcification that is an important part of intervertebral disc degeneration in dogs and people and has been associated with disease in both species. Our findings lead us to speculate that calcification is a process in which initially an amorphous calcium phosphate phase is formed within the nucleus pulposus, and that this then transforms into a crystalline hydroxyapatite. This is not unexpected given what is known about the maturation of bone and teeth and the role of hydroxyapatite in these biological materials. A recent review of the role of calcification in human intervertebral disc disease also points to the role of calcium pyrophosphate ^11^, but we did not identify this specific material in our samples. There may indeed be some differences in the pathological process between dogs and humans, although it is clear that calcium has a significant role in intervertebral disc disease in both species, with canine intervertebral disc disease having been suggested to be a good natural model of the disease in humans ^41^. If our hypothesis regarding the maturation of the calcification in intervertebral discs is correct, it may further be possible to intervene in this process to slow down or even prevent or reverse the process and perhaps reduce the rate of intervertebral disc herniation as a result.

Further investigation into the topographical and mechanical properties of intervertebral disc samples is needed to better describe and verify these findings, as well as attempt to define the biomechanical features of intervertebral disc material.

## Conclusions

Our results shed new light on the process of calcification in intervertebral disc degeneration and disease. It appears that an amorphous calcium phosphate phase is transformed into crystalline hydroxyapatite, and this might either be due to an unidentified trigger or to the loss of a stabilising agent. This change in the mineral phase within the intervertebral disc appears to be related to herniation of the disc, perhaps due to alterations in biomechanical properties. The identification of this process opens up the possibility for further work which might lead to new therapies and/or prophylactic measures to treat and/or reduce the incidence of this frequently disabling disease in both humans and dogs.

## MATERIALS AND METHODS

### Inclusion Criteria

The study population consisted of chondrodystrophic dogs with intervertebral disc extrusion that underwent surgical treatment at two different institutions, the Queen’s Veterinary School Hospital, University of Cambridge and Dick White Referrals, Newmarket, between October 2020 and June 2021. Dogs were included if they had a diagnosis of intervertebral disc extrusion made by either magnetic resonance imaging (MRI) or computed tomography (CT) and confirmed surgically. Details recorded for each case included breed, age, sex and location of extrusion. **The study was approved by the Ethics and Welfare Committee of the Department of Veterinary Medicine, University of Cambridge, and all methods were performed in accordance with the relevant guidelines and regulations. The study is reported in accordance with relevant ARRIVE guidelines. All data is included within submission and as supplementary material**.

### Study and Control Samples

Study samples consisted of extruded disc material collected during decompressive surgery. Material was collected directly from the surgically exposed vertebral canal by physical extraction with a variety of instruments following laminectomy performed with a high-speed burr ^42^.

Control samples consisted of material collected during prophylactic fenestration of adjacent non-extruded discs ^43, 44^. Both extruded and non-extruded disc materials were placed into sterile plain plastic sample pots and held at +4°C for a maximum of 48 hours. They were then transferred and stored at a – 20°C freezer until the freeze-drying process.

If the sample was large enough, half of the collected material was separated for histopathology. These samples were fixed in 10% formalin for 24-48 hours and then transferred into pots with 70% ethanol until histopathologic investigation. For slide preparation, samples were embedded in paraffin wax, cut to thicknesses of 4-5 μm, and stained with haematoxylin and eosin. None of the samples were decalcified during preparation. Slides were then observed with an optical microscope at a magnification of 40x and were evaluated by a European Boarded Anatomic Pathologist. Twenty-seven samples, of which 21 were extruded and 6 were non-extruded, were evaluated in this way, and all were identified as nucleus pulposus because they contained notochordal cells and/or chondrocyte-like cells <u>(data available as supplementary material)</u>.

### Sample Preparation

All samples were freeze-dried prior to mineral analysis for a total of 17 hours. After placing into the freeze-dryer (VirTis^®^ Advantage Plus) they were cooled from +5°C to -35°C at a rate of -0.19°C/min. and at a pressure of 400 mTorr. This was followed by the drying phase where the temperature was gradually increased to +20°C whilst the pressure was held at 80 mTorr. Samples were then brought to room temperature and stored until mineral analysis.

### FTIR Analysis

FTIR analysis was performed with a Shimadzu^®^ IRPrestige21 Fourier Transform Infrared Spectrophotometer equipped with a Golden Gate ATR accessory (Specac Ltd). FTIR scans of samples covered wavenumbers 400 to 4000 cm ^-1^. The spectrometer was set to a scan speed of 2.8 mm/s and had a resolution of 2 cm ^-1^. The ATR sample holder was cleaned twice with 100% ethanol between each sample analysis. Samples were analysed in the form they left the freeze-drying process, with no grinding or further handling / sample prep. Due to the small size of the ATR sample holder and the lack of sample homogeneity most samples (i.e. where possible), were subsampled up to three times and each scan recorded. For samples with multiple scans, when the three spectra differed, the spectrum that had the most intense phosphate peak (900-1200 cm ^-1^) was included in the data analysis.

Additionally, a commercially available synthetic hydroxyapatite powder purchased from Sigma-Aldrich^®^, and an amorphous magnesium calcium phosphate (AMCP) powder synthesised by the Biomineral Research Laboratory, Department of Veterinary Medicine, University of Cambridge, were used as reference minerals during FTIR analysis. These were also analysed with X-ray diffraction to verify that they contained the crystalline materials of the appropriate phase. FTIR spectra of both showed significant PO_4_^-3^ (ν_3_) vibration peaks in the region 1020-1095 cm^-1^. The hydroxyapatite mineral had a sharper PO_4_^-3^ (ν_3_) peak than the AMCP. Moreover, the FTIR spectrum of hydroxyapatite also had an additional PO_4_^-3^ (ν_1_) peak between wavenumbers 955 and 965 cm^-1^ (**Supplementary Figure 2**).

#### Interpretation of FTIR Spectra of IVDs

The extracted FTIR data covered the range between wavenumbers 850 and 1700 cm^-1^, which encompasses the phosphate and carbonate bond vibration regions known to be significant components of calcium phosphate materials ^33, 45, 46^. All FTIR spectra were initially smoothed with a Savitzky-Golay function to filter out high-frequency noise. This, and subsequent, FTIR signal processing was carried out using MATLAB® r2019b (The MathWorks, Inc., see also **Supplementary Figure 4**). Both extruded (study sample) and non-extruded (control sample) spectra were then observed for the visual appearance of the PO_4_^-3^ ν_3_ absorption peak and the presence or absence of the PO_4_^-3^ ν_1_ absorption peak.

Thereafter, second derivative analysis was performed on all the previously smoothed data to obtain the local peak maxima values at key spectral loci, namely: PO_4_^-3^ ν_1_ (955-965 cm^-1^), PO_4_ ^-3^ ν_3_ (1015-1095 cm^-1^) and CO_3_^-2^ ν_3_ (865-890 cm^-1^) stretching modes. R-Studio software (RStudio Team, 2020) was used to compare differences between extruded and non-extruded samples. Before each statistical test/analysis, a Shapiro-Wilk test was performed to confirm that the data were normally distributed.

### X-Ray Diffraction

Powder X-ray diffraction (PXRD) analysis was performed on 5 extruded (study) and 5 non-extruded (control) samples that were previously analysed by FTIR spectroscopy. The freeze-dried disc materials were gently crushed with an agate pestle & mortar and placed on a low background sample holder made of silicone. PXRD data was collected in Bragg-Brentano geometry with a Bruker^®^ D8 Advance diffractometer equipped with a Mo Kα radiation source, and a Lynxeye XE-T linear position sensitive detector with the following parameters: 2θ range: 5-40°; step size 0.015°; time per step 1s; 50 kV x 40 mA.

### Transmission Electron Microscopy Analysis

Transmission Electron Microscopy (TEM) was performed on two representative samples (one extruded and one non-extruded), after they had been freeze-dried and analysed with FTIR spectroscopy and XRD. A Thermofisher Scientific Helios G4 CX scanning electron microscope was used for scanning electron microscopy energy dispersive X-ray (SEM-EDX) analysis, and a Thermofisher Scientific Titan^3^ Themis G2 SuperX 300 kV scanning transmission electron microscope with a Gatan OneView camera was used for TEM, scanning transmission electron microscopy (STEM), energy dispersive X-ray spectroscopy (STEM-EDX) and electron diffraction analysis.

SEM-EDX was initially performed on the extruded sample to determine the elements present in the material before any further sample preparation. A small amount of each of the two freeze-dried disc materials was then suspended in methanol after grinding in a pestle and mortar before being directly drop-cast onto a holey carbon TEM support film (EM Resolutions Ltd.) and air-dried. TEM images of the disc materials were generated with high-angle annular dark-field imaging by scanning transmission electron microscopy (HAADF-STEM). Additionally, electron diffraction patterns were obtained for both samples using a selected area aperture of ∼200 nm diameter at the image plane. Further, STEM-EDX analysis was performed for elemental mapping (data acquired and processed in the Thermo software Velox).

### Raw Data

Raw data may be viewed at the following link: Article Full Data .xlsx - Google Sheets.html

## Supporting information

Appendix containing all supplementary material

## References

1. Bergknut, N., et al. Intervertebral disc degeneration in the dog. part 1: anatomy and physiology of the intervertebral disc and characteristics of intervertebral disc degeneration. Vet. J. 195, 282–291 (2013).

2. Cole, T.C., Burkhardt, D., Frost L., & Ghosh, P. The proteoglycans of the canine intervertebral disc. Biochim. Biophys Acta 839, 127–138 (1985).

3. Cole, T.C., Ghosh, P., & Taylor, T.K.F. Variations of the proteoglycans of the canine intervertebral disc with ageing Biochim. Biophys Acta 880, 209–219 (1986).

4. Hansen, H. A. Pathologic-anatomical study on disc degeneration in dog, with special reference to the so-called enchondrosis intervertebralis. Acta Orthop. Scand. Suppl. 11, 1–117 (1952).

5. Hansen, T., et al. The myth of fibroid degeneration in the canine intervertebral disc: a histopathological comparison of intervertebral disc degeneration in chondrodystrophic and nonchondrodystrophic dogs. Vet. Pathol. 54(6), 945–952 (2017).

6. Rosenblatt, A. J., Bottema, C. D. K., & Hill, P. B. Radiographic scoring for intervertebral disc calcification in the dachshund. Vet. J. 200, 355–361 (2014).

7. Gruber, H., Norton, H. J., Sun, Y., & Hanley, E. N. Crystal deposits in the human intervertebral disc: implications for disc degeneration. J. Spine 7, 444–450 (2007).

8. Karamouzian, S., et al. Frequency of lumbar intervertebral disc calcification and angiogenesis, and their correlation with clinical, surgical, and magnetic resonance imaging findings. J. Spine 35, 881–886 (2010).

9. Shao, J., et al. Differences in calcification and osteogenic potential of herniated discs according to the severity of degeneration based on pfirrmann grade: a cross-sectional study. BMC Musculoskelet. Disord. 17, 191 (2016).

10. Smolders, L. A., et al. Intervertebral disc degeneration in the dog. part 2: chondrodystrophic and non-chondrodystrophic breeds. Vet. J. 195, 292–299 (2013).

11. Zehra, U., et al. Mechanisms and clinical implications of intervertebral disc calcification. Nat. Rev. Rheumatol. 18, 352–362 (2022).

12. Rohdin, C., Jeserevic, J., Viitmaa, R., & Cizinauskas, S. Prevalence of radiographic detectable intervertebral disc calcifications in dachshunds surgically treated for disc extrusion. Acta Vet. Scand. 52, 1–7 (2010).

13. Stigen, Ø., Ciasca, T., & Kolbjørnsen, Ø. Calcification of extruded intervertebral discs in dachshunds: a radiographic, computed tomographic and histopathological study of 25 cases. Acta Vet. Scand. 61(1), 1–9 (2019).

14. Azizaddini, S., Arefanian, S., Redjal, N., Walcott, B. P. & Mollahoseini, R. Adult acute calcific discitis confined to the nucleus pulposus in the cervical spine: case report. J. Neurosurg. Spine 19, 170–173 (2013).

15. Nogueira-Barbosa, M. H., da Silva Herrero, C.F., Pasqualini, W. & Defino, H. L. Calcific discitis in an adult patient with intravertebral migration and spontaneous remission. Skeletal Radiol. 42, 1161–1164 (2013).

16. Rodacki, M. A., Castro, C. E. & Castro, D. S. Diffuse vertebral body edema due to calcified intraspongious disk herniation. Neuroradiol. J. 47, 316–321 (2005).

17. Jin, C., et al. NLRP3 inflammasome plays a critical role in the pathogenesis of hydroxyapatite-associated arthropathy. PNAS USA 108(36), 14867– 14872 (2011).

18. Feinberg, J., Boachie-Adjei, O., Bullough, P. G., & Boskey, A. L. The distribution of calcific deposits in intervertebral discs of the lumbosacral spine. Clin. Orthop. Relat. Res. 254, 303–310 (1990).

19. Lee, R. S., Kayser, M. v., & Ali, S. Y. Calcium phosphate microcrystal deposition in the human intervertebral disc. J. Anat. 208, 13–19 (2006).

20. Taylor, T. K. F., & Little, K. Prolapsed calcified thoracic intervertebral disc. J. Pathol. Bacteriol. 88, 153–157 (1964).

21. Dieppe, P. A., Crocker, P., Huskisson, E. C., & Willoughby, D. A. Apatite deposition disease: a new arthropathy. Lancet 1(7954), 266–269 (1976).

22. Melrose, J., et al. Calcification in the ovine intervertebral disc: a model of hydroxyapatite deposition disease. Eur. Spine J. 18, 479–489 (2009).

23. Henmi, A., et al. Bone matrix calcification during embryonic and postembryonic rat calvarial development assessed by SEM–EDX spectroscopy, XRD, and FTIR spectroscopy. J. Bone Miner. Metab. 34, 41–50 (2016).

24. Lotsari, A., Rajasekharan, A. K., Halvarsson, M., & Andersson, M. Transformation of amorphous calcium phosphate to bone-like apatite. Nat. Commun. (2018).

25. Mahamid, J., Sharir, A., Addadi, L., & Weiner, S. Amorphous calcium phosphate is a major component of the forming fin bones of zebrafish: indications for an amorphous precursor phase. PNAS USA 105(35), 12748–12753 (2008).

26. Nitiputri, K., et al. Nanoanalytical electron microscopy reveals a sequential mineralization process involving carbonate-containing amorphous precursors. ACS Nano 10, 6826–6835 (2016).

27. Brisson B.A., Moffatt S.L., Swayne S.L., & Parent J.M. Recurrence of thoracolumbar intervertebral disk extrusion in chondrodystrophic dogs after surgical decompression with or without prophylactic fenestration: 265 cases (1995-1999). J. Am. Vet. Med. Assoc. 224(11), 1808–1814 (2004).

28. Freeman P. & Jeffery N.D. Re-opening the window on fenestration as a treatment for acute thoracolumbar intervertebral disc herniation in dogs. J. Small Anim. Pract. 58(4), 199–204 (2017).

29. Hellenbrandt, M. The inorganic crystal structure database (ICSD) present and future. Crystallogr. Rev. 10(1), 17–22 (2004).

30. Sudarsanan, K. & Young, R.A. Significant precision in crystal structural details: holly springs hydroxyapatite. Acta Cryst. 25, 1534–1543 (1969).

31. Piazzaa, R. D., et al. Calcium phosphates nanoparticles: the effect of freeze-drying on particle size reduction. Mater. Chem. Phys. 239, (2020).

32. Eisa, M., al Dabbas, M., & Abdulla, F. Quantitative identification of phosphate using x-ray diffraction and fourier transform infrared (FTIR) spectroscopy. Int. J. Curr. Microbiol. Appl. Sci. 4(1), 270–283 (2015).

33. Kourkoumelis, N., & Tzaphlidou, M. Spectroscopic assessment of normal cortical bone: differences in relation to bone site and sex. Sci. World J. 10, 402–412 (2010).

34. Ratner, B. D., Hoffman, A. S., Schoen, F. J. & Lemons, J. E. Biomaterials Science: An Introduction to Materials in Medicine, 2nd Ed. (Elsevier Academic Press Amsterdam 2004).

35. Rey, C., Shimizu, M., Collins, B., & Glimcher, M. J. Resolution-enhanced fourier transform infrared spectroscopy study of the environment of phosphate ion in the early deposits of a solid phase of calcium phosphate in bone and enamel and their evolution with age: 2. investigations in the nu3PO4 domain. Calcif. Tissue In. 49, 383–388 (1991).

36. Miller, L. M., et al. In situ analysis of mineral content and crystallinity in bone using infrared micro-spectroscopy of the v4 PO43-vibration. Biochim. Biophys. Acta 1527, 11–19 (2001).

37. Turunen, M. J., et al. Bone mineral crystal size and organization vary across mature rat bone cortex. J. Struct. Biol. 195, 337–344 (2016).

38. Batcher, K., et al. Phenotypic effects of FGF4 retrogenes on intervertebral disc disease in dogs. Genes 10(6), 435 (2019).

39. Dugat D.R., Bartels K.E, & Payton M.E. Recurrence of disk herniation following percutaneous laser disk ablation in dogs with a history of thoracolumbar intervertebral disk herniation: 303 cases (1994-2011). J. Am. Vet. Med. Assoc. 249(12), 1393–1400 (2016)

40. Grindulis KA, Finlay DB, & Nichol FE. Chemonucleolysis of lumbar intervertebral disc prolapse with chymopapain: outcome after 1 year. Clin. Rheumatol. 6(1), 42–9 (1987).

41. Lee, N.N., et al. Canine models of spine disorders. JOR Spine 3(4), e1109 (2020).

42. Smolders, L. A., et al. Biomechanical assessment of the effects of decompressive surgery in non-chondrodystrophic and chondrodystrophic canine multisegmented lumbar spines. Eur. Spine J. 21, 1692–1699 (2012).

43. Bach, F. C., et al. Potential regenerative treatment strategies for intervertebral disc degeneration in dogs. BMC Vet. Res. 10, 3 (2014).

44. Jeffery, N. D., & Freeman, P. M. The role of fenestration in management of type I thoracolumbar disk degeneration. Vet. Clin. North Am. Small Anim. Pract. 48(1), 187–200 (2018).

45. Chen, K. H., Cheng, W. T., Li, M. J., Yang, D. M., & Lin, S. Y. Calcification of senile cataractous lens determined by fourier transform infrared (FTIR) and Raman microspectroscopies. J. Microsc. 219(1), 36–41 (2005).

46. Gadaleta, S. J., Paschalis, E. P., Betts, F., Mendelsohn, R., & Boskey, A. L. Fourier transform infrared spectroscopy of the solution-mediated conversion of amorphous calcium phosphate to hydroxyapatite: new correlations between x-ray diffraction and infrared data. Calcif. Tissue Int. 58, 9–16 (1996).

