## Appendix containing all supplementary material for "Mineral Phase Changes During Intervertebral Disc Degeneration"

### Supplementary Figure 1

Clinical Data of Patients from which Intervertebral Disc Materials were Collected

| | | | |
|---|---|---|---|
| Breed<br>(%, n) | Dachshund | 38.6 % | 22 |
| | French Bulldog | 24.6 % | 14 |
| | Cocker Spaniel | 15.8 % | 9 |
| | Mixed Breed | 10.5 % | 6 |
| | Shih Tzu | 3.5 % | 2 |
| | Beagle | 1.7 % | 1 |
| | Cavalier King Charles Spaniel | 1.7 % | 1 |
| | Chihuahua | 1.7 % | 1 |
| | Maltese | 1.7 % | 1 |
| Age<br>(months) | $\bar{x}$ | 70.8 | |
| | Median | 66 | |
| | SD | 28.5 | |
| | Max. | 156 | |
| | Min. | 30 | |
| Gender<br>(%, n) | M | 14% | 8 |
| | MN | 63.1% | 36 |
| | F | 5.2% | 3 |
| | FN | 17.5% | 10 |
| Weight<br>(kg) | $\bar{x}$ | 10.3 | |
| | Median | 8.9 | |
| | SD | 4.6 | |
| | Max. | 19.5 | |
| | Min. | 2.7 | |
| Time between date of sample collection and date of onset<br>(days) | $\bar{x}$ | 14 | |
| | Median | 2 | |
| | SD | 29.8 | |
| | Max. | 171 | |
| | Min. | 0 | |
| Localisation of the extruded IVD<br>(%, n) | Cervical | 15.5 % | 9 |
| | Thoracolumbar | 84.5% | 49 |
| Clinical Score<br>(%, n) | Grade 1 | 12.7% | 7 |
| | Grade 2 | 35% | 20 |
| | Grade 3 | 35% | 20 |
| | Grade 4 | 7% | 4 |
| | Grade 5 | 10.5% | 6 |
| Pain on Palpation<br>(%, n) | Absent Pain | 12.3% | 7 |
| | Mild Pain | 52.6% | 30 |
| | Significant Pain | 35% | 20 |

### Supplementary Figure 2

Second derivative FTIR spectra of test materials

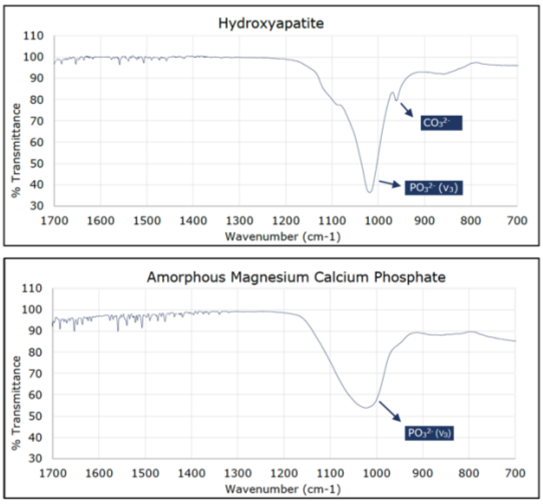


### Supplementary Figure 3

*Second derivative FTIR Spectra (850-1700 cm$^{-1}$) of  Extruded IVD Materials Tested*

*Second derivative FTIR Spectra (850-1700 cm$^{-1}$) of Non-extruded IVD Materials Tested*

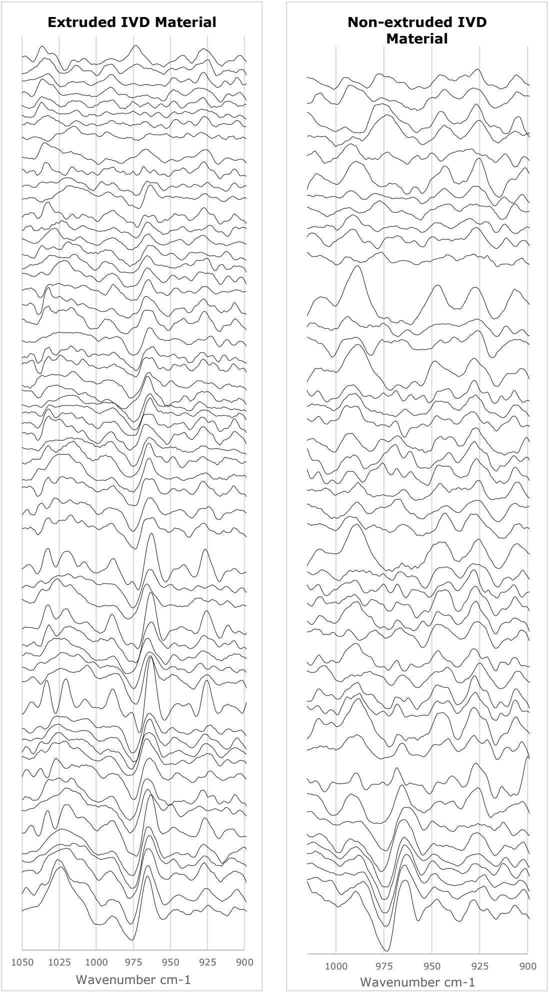

### Supplementary Figure 4

*MATLAB Codes: FTIR Signal Processing Script*

**Data Import -** clear the memory and command window

```
clear
close all
clc
format long

% import datafile details
datafiles = dir('**/*.txt');
index = contains({datafiles.name}, 'smooth');
datafiles(index)=[];
clear index

% format filenames for import
filenames = {datafiles(:).name};
filenames = filenames';
filenames = strrep(filenames,'.txt','');

% Batch import of FTIR data into a single data matrix
% comprising wavelength (:,1,:) and transmission (:,2,:) values for each measurement file
(:,:,i)
for i = 1:numel(datafiles(:,1))
   name = char(filenames(i,:));
   data(:,:,i) = readmatrix(name);
end

% Remove NaN columns
for i = 1:numel(datafiles(:,1))
dat(:,:,i) = data(:,~all(isnan(data(:,:,1))),i);
end
data = dat; clear dat
```

**Data Smoothing -** 2nd order Savitzky-Golay filter applied to transmission values using a frame-length of 15 smoothed transmission values appended to data matrix as a new column (:,3,:)

```
datacols = size(data,2);
for i = 1:numel(datafiles(:,1))
   data(:,datacols+1,i) = sgolayfilt(data(:,2,i),2,15);
end

clear i datacols
```

**Export Data -** export of the smoothed transmission data

```
for i = 1:numel(datafiles(:,1))
  txt1(i) = strcat(string(filenames(i)),'-smoothed.txt');
  writematrix(data(:,:,i),txt1(i));
end

clear i
```

**Plot and Save Smoothed spectra**

```
for i = 1:numel(datafiles(:,1))
    figure
    plot(data(:,1,i),data(:,3,i))
    xlabel('Wavenumber/cm^{-1}')
    ylabel('Transmittance')
    xlim([500 4000])
    ylim([30 130])
    box off;
    set(gca,'TickLength',[0.015 0.015]);
    set(gca,'TickDir','out');
    set(gca,'LineWidth',1);
    set(gca,'xdir','reverse');
    txt2(i) = strcat(string(filenames(i)),'-smoothed');
    an1 = annotation('textbox',[0.15, 0.12, 0.1,
0.1],'string',txt2(i),'FitBoxToText','on','EdgeColor','white');
    export_fig(txt2(i),'-tif','-q101')
end
```

**Taking and Smoothing Derivatives**

```
    clear dat idx xfilt xcols a i k n p q w x;

    r = [1:numel(data(1,1,:))];

% Differential parameters
    n = 2;                            % nth derivative

% Smoothing parameters
    passes = 3;                       % passes of boxcar filter
    windowSize = 5;                   % window size(s)
    a = 1;                            % numerator

% Plotting parameters
    X_MIN = 850;                      % Axis limits
    X_MAX = 1150;

% Differentiation
 for i = 1:numel(r)
  x(:,:,i) = diff(data(:,2,i),n,1);
 end
```

**Plotting and export of derivative spectra after several passes of box-car filter**

```
b = (1/windowSize)*ones(1,windowSize);

for i = 1:numel(x(1,1,:))
    for p = 1:passes
        x(:,p+1,i) = filter(b,a,x(:,p,i),[],1);
    end
    figure
    plot(data(2:numel(x(:,1,i))+1,1,i),x(:,passes+1,i))
    txt2(i) = strcat(string(filenames(i)),'-derivative');
    an1 = annotation('textbox',[0.15, 0.12, 0.1,
0.1],'string',txt2(i),'FitBoxToText','on','EdgeColor','white');
```

```
    xlim([X_MIN X_MAX]);
    ylim([-0.1 0.1]);
    xlabel('Wavenumber/cm^{-1}')
    ylabel('Second Derivative')
    set(gca,'xdir','reverse');
    set(gca,'TickLength',[0.015 0.015]);
    set(gca,'TickDir','out');
    set(gca,'LineWidth',1);
    %export_fig(txt2(i),'-tif','-q101')
end
```

**Export Derivative Data**

```
for i = 1:numel(datafiles(:,1))
  txt1(i) = strcat(string(filenames(i)),'-derivative.txt');
  dat(:,1) = data(2:numel(x(:,1,i))+1,1,i);
  dat(:,2) = x(:,passes+1,i);
  writematrix(dat,txt1(i));
end
clear i
```

**Derivative Spectra Local Maxima**

```
% capture the magnitude of local maxima at specific wavelength ranges for
% each measurement file

% find max at 960 cm-1

clear g i d ix diff_index waves diff_peak_range diff_peak_vector diff_peaks

diff_index(1,1) = 955;
diff_index(2,1) = 965;

for g = 1:numel(data(1,1,:))
    waves = data(:,1,g)';
    for i = 1:2
        [d, ix] = min(abs(waves - diff_index(i,1)));
        diff_peak_range(i,1) = ix;
    end
    diff_peak_vector = x(diff_peak_range(1):diff_peak_range(2),passes+1,g);
    diff_peaks(g,1) = max(diff_peak_vector);
end

% find max at 880 cm-1

diff_index(1,1) = 870;
diff_index(2,1) = 890;

for g = 1:numel(data(1,1,:))
    waves = data(:,1,g)';
    for i = 1:2
        [d, ix] = min(abs(waves - diff_index(i,1)));
        diff_peak_range(i,1) = ix;
    end
    diff_peak_vector = x(diff_peak_range(1):diff_peak_range(2),passes+1,g);
    diff_peaks(g,2) = max(diff_peak_vector);
end
```

```
% find max at 1030 cm-1

diff_index(1,1) = 1025;
diff_index(2,1) = 1041;

for g = 1:numel(data(1,1,:))
    waves = data(:,1,g)';
    for i = 1:2
        [d, ix] = min(abs(waves - diff_index(i,1)));
        diff_peak_range(i,1) = ix;
    end
    diff_peak_vector = x(diff_peak_range(1):diff_peak_range(2),passes+1,g);
    diff_peaks(g,3) = max(diff_peak_vector);
end

% find max between 850-1150 cm-1

diff_index(1,1) = 850;
diff_index(2,1) = 1150;

for g = 1:numel(data(1,1,:))
    waves = data(:,1,g)';
    for i = 1:2
        [d, ix] = min(abs(waves - diff_index(i,1)));
        diff_peak_range(i,1) = ix;
    end
    diff_peak_vector = x(diff_peak_range(1):diff_peak_range(2),passes+1,g);
    diff_peaks(g,4) = max(diff_peak_vector);
end

% find max between 1520-1660 cm-1

diff_index(1,1) = 1520;
diff_index(2,1) = 1660;

for g = 1:numel(data(1,1,:))
    waves = data(:,1,g)';
    for i = 1:2
        [d, ix] = min(abs(waves - diff_index(i,1)));
        diff_peak_range(i,1) = ix;
    end
    diff_peak_vector = x(diff_peak_range(1):diff_peak_range(2),passes+1,g);
    diff_peaks(g,5) = max(diff_peak_vector);
end
```

**Supplementary Figure 5**

*STEM-EDX elemental mapping of the Extruded IVD Sample*

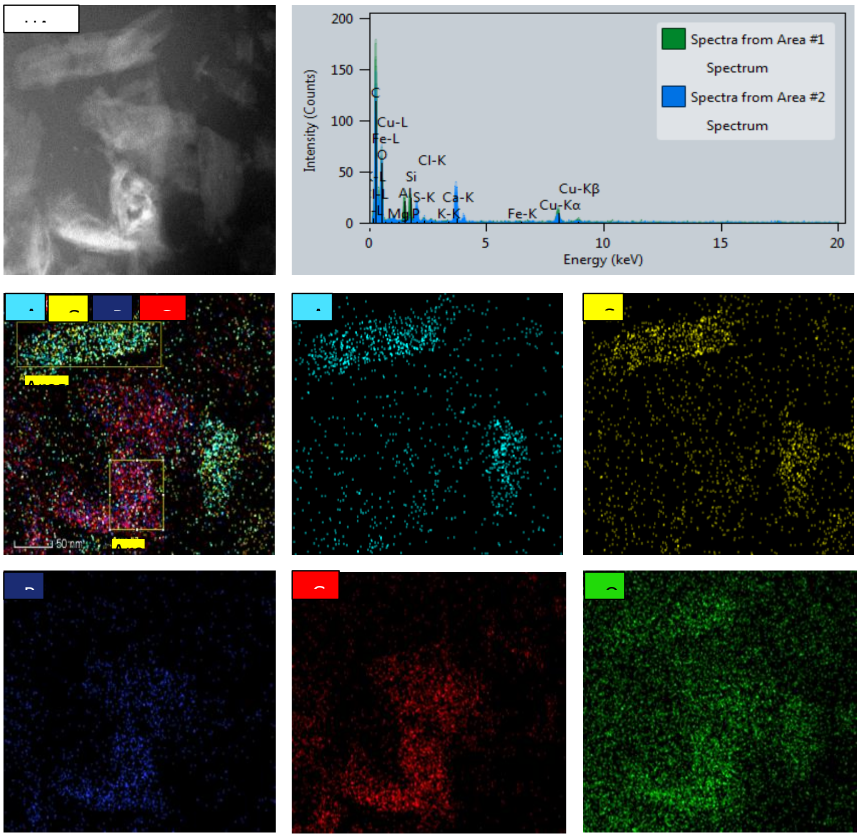


*Note.* (Area 1) the blocky structure on the TEM image are Al, Si and O rich; (Area 2) the needle-like structures are Ca, P and O rich

**Supplementary Figure 6**

*TEM and HAADF-STEM Images of IVD Material*

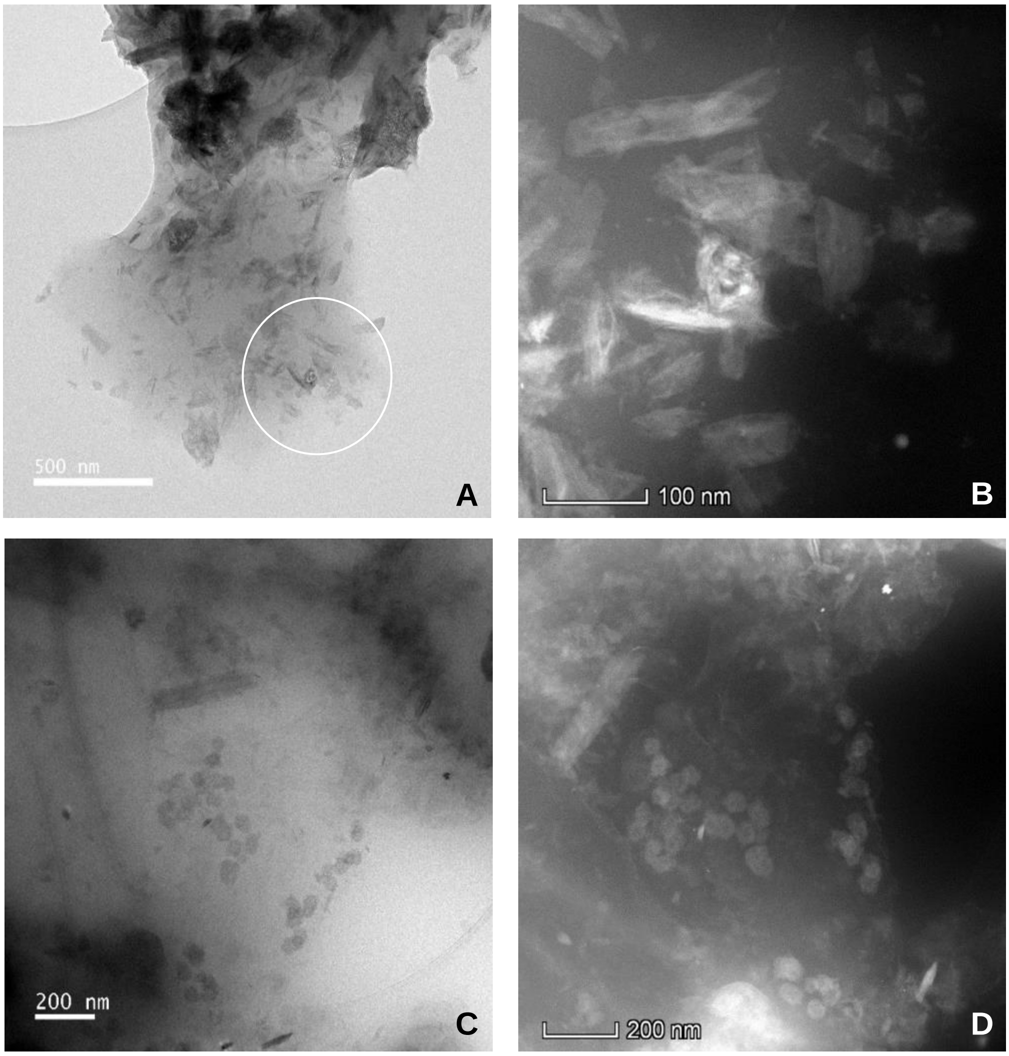


*Note*. (A) Extruded IVD Material: TEM image revealing particles in gel-like substance; (B) Extruded IVD Material: HAADF-STEM images of the circled area on image A showing some 'blocky' particles and some needle agglomerates; (C) Non-extruded IVD Material: TEM shows blocks and spheres in gel; (D) Non-extruded IVD Material: HAADF-STEM image confirmative of the TEM the image.

### Supplementary Figure 7

*STEM-EDX Elemental Mapping of the Non-extruded IVD Sample*

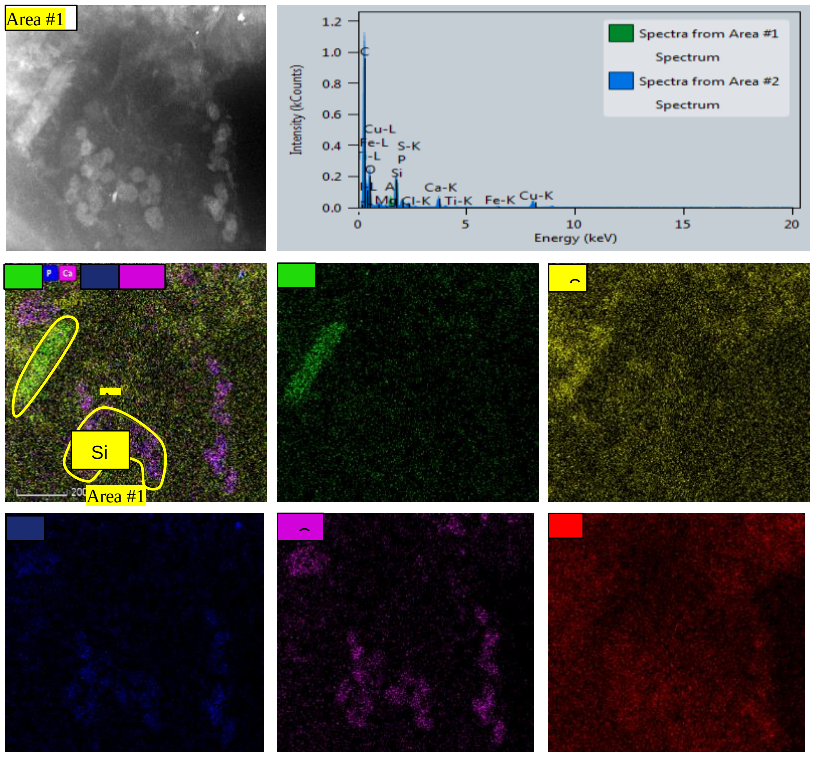


*Note*. (Area 1) the blocky structure on the TEM image are Al, Si and O rich; (Area 2) the amorphous spheres mainly contain Ca, P and O.

### PXRD data analysis

An empirical background collected on a blank silicon low background sample holder was first subtracted from all diffractograms. Rietveld analysis was performed with program TOPAS Academic V6.[1] The structure of hydroxylapatite was retrieved from the Inorganic Crystal Structure Database.[2] No structural parameters but unit cell, one isotropic thermal parameter for all atoms, calcium and hydroxyl ions occupancy factors, were refined. The March-Dollase model for preferred orientation[3] was applied on the crystallographic planes parallel to (0 0 1). A shifted Chebyshev function with ten parameters was used to fit the background. A $LaB_6$ NIST standard[4] was used to model the instrumental fundamental parameters. Hydroxylapatite contribution to peak broadening for crystallite size was modelled isotropically using a Lorentzian function with full width half maximum depending on $2\theta$ according to the Scherrer equation. A representative Rietveld refinement plot is shown in Supplementary Figure 8. Supplementary Table 1 reports relevant output parameters from the Rietveld refinements.

### Supplementary Figure 8

*Experimental (black dots), calculated (red line) and difference (grey line) patterns for the Rietveld refinement of sample SN4002-1. Peak position marks for HA are shown in blue.*

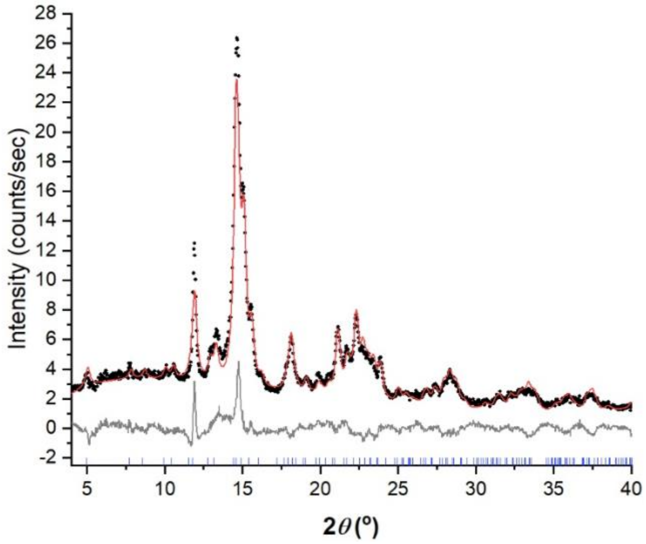


### Supplementary Table 1

*Rietveld refinements output parameters for all crystalline samples analysed.*

| | $R_{wp}$ (%) | $X^2$ | *a* (Å) | *c* (Å) | Occ(Ca1) | Occ(Ca2) | Occ(OH) | Cr.Size (nm) |
|---|---|---|---|---|---|---|---|---|
| **SN1010** | 11.23 | 0.13 | 9.42(1) | 6.887(7) | 0.27(1) | 1.00(3) | 0.49(6) | 12.9(8) |
| **SN1011** | 9.88 | 0.19 | 9.437(4) | 6.901(3) | 0.303(5) | 1.000(9) | 0.58(2) | 14.3(4) |
| **SN1029** | 8.32 | 0.12 | 9.42(1) | 6.901(8) | 0.22(1) | 1.00(4) | 0.37(7) | 12.8(8) |
| **SN4002** | 14.74 | 0.13 | 9.42(9) | 6.88(1) | 0.28(3) | 1.00(7) | 0.49(1) | 13(2) |
| **SN4005** | 6.14 | 0.10 | 9.46(1) | 6.907(1) | 0.256(7) | 0.92(2) | 0.17(5) | 7.5(3) |

### References


[1] Coelho, A.A., 2018. TOPAS and TOPAS-Academic: an optimization program integrating computer algebra and crystallographic objects written in C++. Journal of Applied Crystallography, 51(1), pp.210-218.

[2] Hellenbrandt, M., 2004. The inorganic crystal structure database (ICSD)—present and future. Crystallography Reviews, 10(1), pp.17-22.
[3] Dollase, W.A., 1986. Correction of intensities for preferred orientation in powder diffractometry: application of the March model. Journal of Applied Crystallography, 19(4), pp.267-272.
[4] Black, D.R., Windover, D., Henins, A., Filliben, J. and Cline, J.P., 2011. Certification of standard reference material 660B. Powder Diffraction, 26(2), pp.155-158.
